# IDENTIFICATION OF A NOVEL MECHANISM FOR LFA-1 ORGANIZATION DURING NK CYTOLYTIC RESPONSE

**DOI:** 10.1101/2020.10.27.357970

**Authors:** Alejandro P. Pariani, Evangelina Almada, Florencia Hidalgo, Carla Borini-Etichetti, Rodrigo Vena, Leandra Marín, Cristián Favre, James R. Goldenring, M. Cecilia Larocca

## Abstract

The elimination of transformed and viral infected cells by natural killer (NK) cells requires the formation of a specialized junction between NK and target cells, generally denominated immunological synapse (IS). After initial recognition, the IS enables the directed secretion of lytic granules content into the susceptible target cell. The lymphocyte function–associated antigen (LFA)-1 regulates NK effector function by enabling NK-IS assembly and maturation. The pathways underlying LFA-1 accumulation at the IS in NK cells have not been elucidated so far. A kinase anchoring protein 350 (AKAP350) is a centrosome/Golgi associated protein with a central role in the regulation of Golgi organization and microtubule dynamics. In T cells, AKAP350 participates in LFA-1 activation by mechanisms that remain elusive. We first investigated AKAP350 participation in NK cytolytic activity. Our results showed that the decrease in AKAP350 levels by expression of specific shRNAs (AKAP350KD) inhibited NK-YTS cytolytic activity, without affecting NK:target cell conjugate formation. The impairment of NK effector function in AKAP350KD cells correlated with decreased LFA-1 clustering and defective IS maturation. AKAP350KD cells that were exclusively activated *via* LFA-1 showed impaired LFA-1 organization and deficient lytic granule translocation as well. In NK AKAP350KD cells, activation signaling through Vav1 was preserved up to 10 min of interaction with target cells, but significantly decreased afterwards. Experiments in YTS and in *ex vivo* NK cells revealed the existence of an intracellular pool of LFA-1 vesicles, which partially associated with the Golgi apparatus. Upon NK cell activation, this LFA-1 pool redistributed to the IS in a Golgi-AKAP350-dependent manner. Disruption of Golgi integrity or alteration of microtubule dynamics impaired LFA-1 localization at the IS. Therefore, this study reveals the existence of a Golgi/AKAP350 dependent trafficking pathway for LFA-1, which is relevant for LFA-1 organization at NK-lytic IS and, concomitantly, for NK cytolytic function.

## 1 INTRODUCTION

Natural killer (NK) cells are the cytolytic effectors of the innate immune system, which constitute the first line of immunological defense and are particularly relevant for the elimination of viral infected and neoplastic cells. NK cytotoxic effector function relies on the formation of a specialized junction with target cells, generally referred to as immunological synapse (IS). The formation and maturation of NK-IS requires the participation of multiple NK receptors, signaling molecules and cytoskeletal elements, which accumulate in specific regions within the IS to form a supramolecular activation cluster (SMAC), distinctly organized at peripheral (pSMAC) and central (cSMAC) zones (Orange, 2008). The persistent outside-in signaling generated by the interaction of NK activating receptors with their ligands in the target cell at the IS, in a context of absence of inhibitory signals, triggers a multistep, highly regulated process that culminates with the directed secretion of lytic molecules contained in specialized lysosomes (lytic granules) through the cSMAC into the target cell (Rak et al., 2011). The initial contact between NK and target cells may involve any of a number of different receptors, among which the integrin leukocyte functional antigen (LFA)-1 is of particular interest. LFA-1 is a heterodimer consisting of αL (CD11a) and β2 (CD18) chains, which is essential for the stabilization of the IS (Hoffmann et al., 2011). In addition to mediating the tight adhesion to target cells, LFA-1 contributes to NK cytotoxic response by initiating signaling pathways that promote lytic granule convergence to the centrosome and lytic granule/centrosome polarization towards the IS (Barber et al., 2004; Bryceson et al., 2005). LFA-1 reorganization implies conformational changes that lead to an activated configuration, as well as an increase in its localization at the NK-target cell contact site. The implication of different NK receptors and interleukins in the inside out signaling that can lead to LFA-1 activation has been characterized (Urlaub et al., 2017). Nevertheless, the cellular mechanisms underlying the increase in LFA-1 expression at the IS in this context remain unclear.

A major mediator of NK cell cytolytic activity is the cytoskeleton. In this regard, the role of actin dynamics on the maturation of the lytic IS has been well established (Lagrue et al., 2013). Regarding the microtubule cytoskeleton, it remains unclear whether they serve any role other than facilitating the delivery of lytic granules to the synaptic cleft (Chen et al., 2006; Stinchcombe et al., 2006). The A-kinase anchoring protein 350 (AKAP350), also known as AKAP450, CG-NAP or AKAP9, is a centrosome and Golgi-scaffold protein (Schmidt et al., 1999) that regulates the Golgi apparatus organization (Larocca et al., 2004) and has a prominent role in microtubule remodeling (Takahashi et al., 2002; Larocca et al., 2006; Rivero et al., 2009). The first study that revealed AKAP350 participation in the regulation of immune cells function was performed in migratory T cells (El Din El Homasany et al., 2005). Those studies demonstrated that the interference with AKAP350 function at the centrosome leads to inhibition of T cell migration on cells coated with LFA-1 ligand ICAM-1. AKAP350 participation in LFA-1 induced cell migration was further confirmed by later studies that showed that AKAP350 depletion leads to defective T-cell migration on ICAM-1 coated surfaces (Ong et al., 2018). An independent study indicated that, during antigen recognition by T-cells, AKAP350 participates in T cell receptor and LFA-1 activation (Robles-Valero et al., 2010). Regarding the role of AKAP350 in LFA-1 activation in NK cells, which have very different requirements and implications from those in T cells (Watz and Long, 2010), they have not been addressed so far.

In the present study we analyzed AKAP350 participation in the development of NK cytotoxic effector function and characterized its role in LFA-1 reorganization at the lytic IS. Our results reveal a novel role for the Golgi apparatus in LFA-1 compartmentalization and, concomitantly, in NK lytic IS maturation.

## 2 MATERIALS AND METHODS

### 2.1 NK and target cell lines

The immortalized NK YTS cells (Drexler and Matsuo, 2000) and KT86 cells, derived from the MHC class I-negative K562 erythroleukemia cell line stably expressing CD86 (Parry et al., 2003) were obtained from Dr. Jordan Orange lab. Cells were maintained in RPMI medium 1640 supplemented with 10% FBS, 1% L-glutamine, 1% non-essentials amino acids and 1% streptomycin/ampicillin mixture at 37°C and 5% CO_2_ atmosphere.

In order to prepare YTS cells with reduced expression of AKAP350 (AKAP350KD), we proceeded as described (Tonucci et al., 2018). Details are included in Supplementary Methods. To delocalize AKAP350 from the Golgi apparatus, the nucleotide sequence codifying for AKAP350(1-1229) (AKAP350GBD) was cloned into pInducer20 vector (details at http://www.addgene.org). Lentiviral particles were constructed as described in Supplementary Methods. Transduced YTS cells were subjected to G418 selection (1 μg/ml) for two weeks. AKAP350GBD expression was induced with Doxycycline (2 μg/ml) 24 h before the experiments.

*Ex vivo* NK cell purification and culture. Blood samples from volunteer donors were used to prepare *ex vivo* NK cells. All human samples were obtained after informed patient consent and were used with approval of the institutional internal review board for the protection of human subjects of the Hospital Provincial del Centenario (Rosario, Argentina). NK cells were purified by negative selection using an RosetteSep Human NK cells Enrichment cocktail (StemCell # 15025) and cultured in IL-2 supplemented medium as detailed in Supplementary Methods. *Ex vivo* NK cells with reduced expression of AKAP350 were prepared by transduction with shRNA expression lentiviral particles as described above.

### 2.2 CFSE-based Cytotoxic assay

In order to determine the lytic capacity of YTS cells, we proceed as described (Jedema et al., 2004). KT86 cells were washed with PBS, resuspended at 2 x10^6^ cells/ml and labeled with CFSE (300 nM, BD Horizont) for 15 min at 37°C. The reaction was stopped by the addition of an equal volume of fetal bovine serum (FCS). CFSE-labeled target cells were washed and resuspended in RPMI complete medium. Cell concentration was adjusted to 5 × 10^5^ cells/ml, and 100 μl/tube were plated in Eppendorf tubes. YTS cells were added at 2:1 or 10:1 effector:target ratios. Tubes were incubated in a humidified atmosphere of 5% CO2 and 37°C for 3 hours. Dead cells were stained with propidium iodide (1 μg/mL) and samples were directly analyzed by flow cytometry using a BD FACSAria II flow cytometer. 10000 events were acquired for each sample. Double positive events for CFSE and PI were considered dead target cells. The percentage of specific lysis was calculated as follows: specific lysis (%) = (Double positive cells / CFSE positive cells) x 100.

### 2.3 FACS-based conjugation assay

Target cells were labeled with CFSE as described above. YTS cells were washed with PBS, resuspended to 1 × 10^6^ cells/ml and labeled with Cell TrackerTM Deep Red (1.5 mM, Invitrogen) for 20 min at 37°C. Cells were combined at different YTS:KT86 ratios, incubated and fixed, and conjugate formation analyzed by flow cytometry as detailed in Supplementary Methods.

### 2.4 Immunofluorescence

KT86 were labeled with CFSE as described above. Conjugates between YTS cells and KT86 at a 2:1 ratio were established in suspension for 15 min at 37°C and adhered to poly-lysine-coated glass slides (Polyprep; Sigma-Aldrich) for additional 15 min. In experiments evaluating NK cell activation by specific activating receptors, poly-l-lysine coated glass slides were coated by overnight incubation with ICAM-1 (5 μg/mL, Biolegend 552906) or with ICAM-1 and anti-CD28 (5 μg/mL, BDbioscience 554121) in PBS. Slides were washed, and NK cells were incubated on the slide for 30 min at 37°C. Cells were washed, fixed and stained, and images were acquired by confocal microscopy as detailed in Supplementary Methods.

### 2.5 Image analysis

Lytic granule convergence and centrosome and lytic granule polarization towards the IS. YTS:KT86 conjugates were stained with anti γ-tubulin, perforin and DAPI, for lytic granules, centrosome and nucleus identification, respectively. The IS was defined as the cell to cell contact region identified in the images obtained by differential interference microscopy (DIC). The cell perimeter of the YTS cell forming the IS was drawn using the same images. At the *x,y* plane corresponding to the centrosome center, a threshold was set on the γ-tubulin or perforin channel to define the respective masks, which were used to automatically outline the centrosome or the lytic granule regions. The *x,y* coordinate values for the IS, the lytic granules, the centrosome, and the cell centroids were determined using the appropriate ImageJ tool and the distances from the lytic granules, the centrosome or the cell centroids to the IS centroid, or from the lytic granules to the centrosome centroid were calculated. In the case of lytic granules, an average area-weighted distance (AWD) was calculated for each cell by applying a modified use of Shepard’s Method (Mentlik et al., 2010):

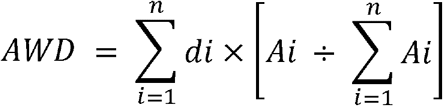

where ***Ai*** is the area of each particular lytic granule region and ***di*** is the distance from this particle to the IS or the centrosome. This modification allowed weighting the distances by the area of each lytic granule region, which is important considering that clusters of lytic granules could inappropriately be discerned as an individual granule of a larger area and, therefore, could be under-represented if a factor considering each granule area is not used. For analysis of lytic granules and centrosome translocation during specific LFA-1 mediated activation, the surface of YTS interaction with ICAM-1 coated slides was considered as the IS site, and the distance was measured at the *y,z* plane.

LFA-1-vesicles polarization towards the IS. YTS:KT86 or *ex vivo* NK:KT86 cell conjugates were permeabilized and stained with anti LFA-1. The IS and the cell regions were defined as explained above. A threshold was set on the LFA-1 channel to define a mask, which was used to automatically outline the regions corresponding to LFA-1 vesicles. The IS, LFA-1 vesicles and the cell centroids were determined and the AWD or the distances from LFA-1 vesicles or from the cell centroids to the IS centroid were calculated similarly to what we explained above for lytic granules.

LFA-1 accumulation and organization at the IS. YTS or *ex vivo* NK cells conjugated with KT86 cells were analyzed. DIC images were used for IS and cell selections. For Figure 3a, the area of the LFA-1 positive region at the IS was additionally determined in the LFA-1 channel. Average and total intensity of fluorescence was measured at NK-IS and at the whole cell and the percentage of protein accumulation at the IS were calculated. The image J tool “reslice” was used for *x,z* planes reconstruction of the IS region and the plugin Radial Profile (http://rsbweb.nih.gov/ij/plugins/radial-profile.html) was used for measuring the intensity of fluorescence at concentric circumferences beginning at the center towards the periphery of the IS.

Centrosomal localization of AKAP350. Masks at γ-tubulin and AKAP350 channels were defined to automatically outline centrosomal and AKAP350 total pixels. Total intensity of fluorescence in the AKAP350 channel was measured in each region of interest, and the percentage of AKAP350 at the centrosomes was calculated for each cell.

AKAP350 or LFA-1 association to the Golgi apparatus. A threshold was set in the GM130 channel to define a mask, which was used to automatically outline the Golgi pixels, and the cell region was delimited in the DIC channel. Total intensity of fluorescence in the AKAP350 or LFA-1 channel was measured in each region of interest, and the percentage of AKAP350 or LFA-1 fluorescence at the Golgi apparatus was calculated for each cell. A similar protocol was used to estimate LFA-1 association with the recycling endosomes, using Rab11 staining to define this compartment.

### 2.6 Immunoblotting

Cells were harvested at 400 *g* for 5 min at room temperature and washed with cold PBS. Pelleted cells were lysed and western blot analysis performed as we have previously described (18). Specific details are included in Supplemmentary Methods. For signaling analysis, we used the protocol described by Robles-Valero *et al*. (2010) adapted to NK cells. YTS cells and KT86 were mixed at a 6:1 effector:target ratio and centrifuged for 15 seconds at 100 *g* to facilitate conjugate formation. Cells were incubated at 37° C and 5% CO for different periods. The cells were harvested and lysed in ice-cold lysis buffer with phosphatase inhibitors.

### 2.7 YTS cells treatments

In order to assess the role of microtubules on IS formation, YTS cells were harvested and 2×10^6^ cells were treated for 1 hour with nocodazole (17 μM) on ice or for 1 hour with Taxol (10 μM) at room temperature and then allowed to form conjugates with KT86 cells at a 2:1 ratio for 30 min at 37°C. In order to evaluate the role of the Golgi apparatus in LFA-1 organization during IS formation, YTS cells were treated with Brefeldin A (5 μg/ml) for 30 min at 37°C. After Brefeldin A treatment, YTS cells were allowed to form conjugates with KT86 cells at a 2:1 ratio for 30 min at 37°C. YTS cells without treatment were used as controls.

### 2.8 Statistical analysis

Data are expressed as mean ± s.e.m. Paired Student’s t-test was used for comparison between groups and non-parametric Mann–Whitney test was used for comparisons within each experiment. p < 0.05 was considered statistically significant.

## 3 RESULTS

### 3.1 Decrease of endogenous AKAP350 expression impairs NK cytolytic activity by impairing IS maturation

We first evaluated AKAP350 expression in YTS immortalized NK cells. Western blot analysis of YTS cell-extracts confirmed AKAP350 expression at protein level. Immunofluorescence analysis of AKAP350 staining in resting YTS cells showed that AKAP350 colocalized with γ-tubulin and GM-130, what is indicative of AKAP350 localization at the centrosome and at the Golgi apparatus (Supplementary Figure S1).

To analyze the impact of AKAP350 expression in the development of NK cytolytic effector function, we generated YTS cells with decreased AKAP350 expression (AKAP350KD) using a shRNA expression system for targeting two different AKAP350 mRNA sequences (shRNA1 and shRNA4). Western blot analysis of AKAP350 protein levels showed that the expression of either shRNA1 or shRNA4 efficiently decreased AKAP350 levels to less than 10% of control levels (Figure 1a). To assess AKAP350 participation in the formation of NK:target cell conjugates, control or AKAP350KD YTS cells were labeled with different fluorophores and incubated at 2:1 or 10:1 NK:target cell ratios for different periods. The formation of cell conjugates positive for both fluorophores was analyzed by flow cytometry. Our results showed that the decrease in AKAP350 expression did not elicit any evident effect in the initial adherence between NK and their susceptible target cells (Supplementary Figure S2). In order to analyze NK cytolytic activity in those conditions, control and AKAP350KD YTS cells were incubated with CFSE-labeled KT86 cells. Target cell lysis was analyzed by flow cytometry using propidium iodide to identify dead cells (Figure 1b, c). Our results showed that the decrease in AKAP350 levels by expression of either shRNA1 or shRNA4 impaired NK cytolytic activity.

**Figure 1.**
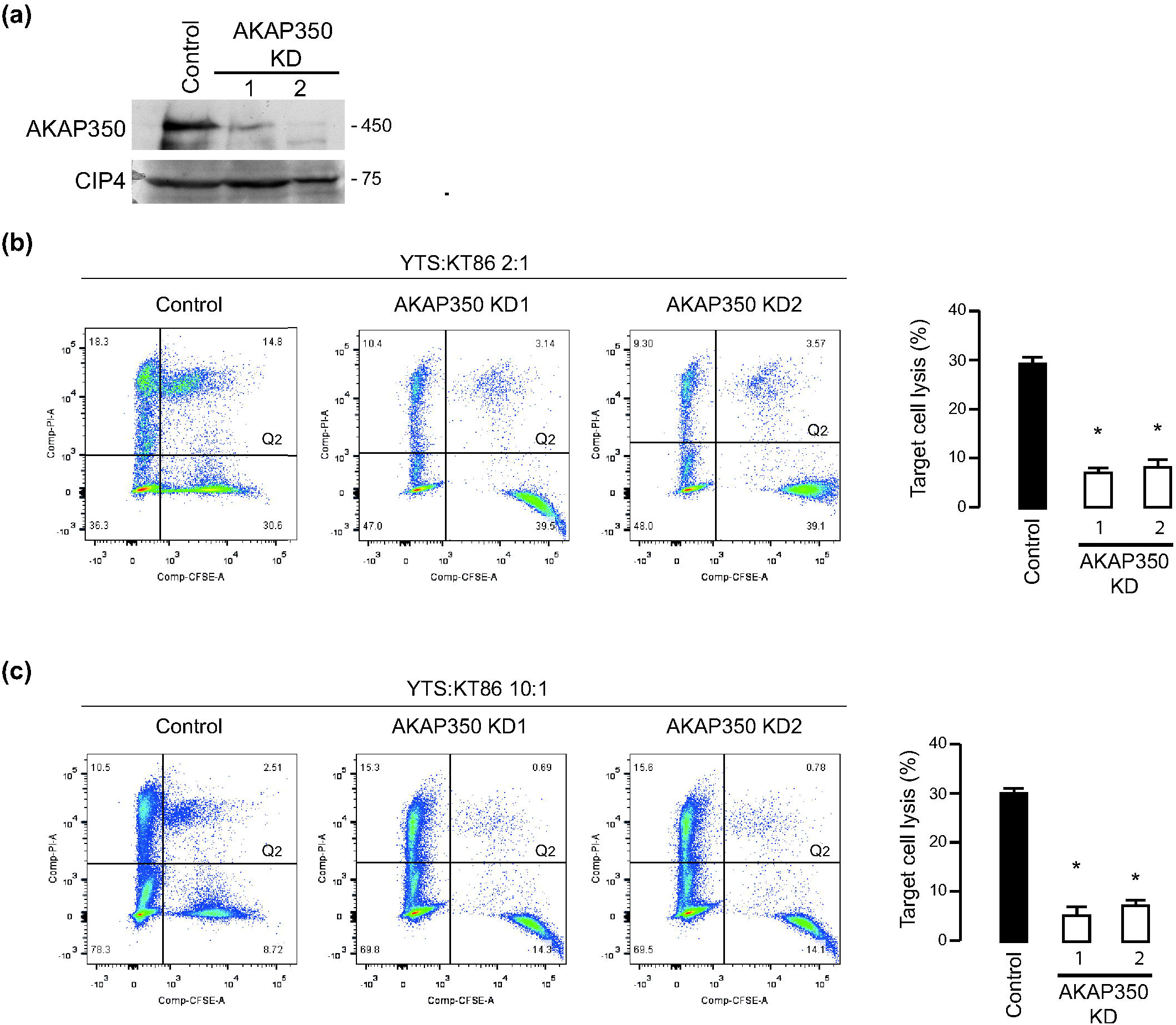
Reduction of AKAP350 expression levels diminishes YTS cells’ cytotoxicity. Two different specific shRNAs (shRNA1 and shRNA4) were used to generate YTS cells with decreased AKAP350 expression (AKAP350KD1 and AKAP350KD2, respectively). **(a)** Western blot analysis of AKAP350 expression in control, AKAP350KD1 and AKAP350KD2 YTS cells. CDC42-interacting protein 4 (CIP4) was used as loading control. **(b,c)** KT86 cells were stained with CFSE and mixed with YTS cells at effector: target ratios of 2:1 (b) or 10:1 (c). After the incubation period, cells were stained with PI and analyzed by flow cytometry. YTS cytotoxic activity was estimated as the fraction of total CFSE positive events that were also positive for PI (Q_2_). Bars chart represents the mean percentage of double positive events for control, AKAP350KD1 and AKAP350KD2 cells, representative of three independent experiments. *p < 0.05

Considering that lytic granule translocation towards the IS is an essential step in NK cytolytic response, lytic granule localization was evaluated in control and AKAP350KD YTS cells conjugated to CFSE-labeled KT86 target cells. Perforin and γ-tubulin staining were used to identify lytic granules and the centrosome, respectively. The average distance between lytic granules and the IS was increased in AKAP350KD cells, thus indicating decreased lytic granule translocation to the IS in these cells (Figure 2a). Regarding the steps that lead to lytic granule translocation to the IS, both lytic granule convergence to the centrosome and centrosome translocation towards the IS were impaired in AKAP350KD cells, as denoted by the increased distance from the lytic granules to the centrosome (Figure 2a) and from the centrosome to the IS (Figure 2b).

**Figure 2.**
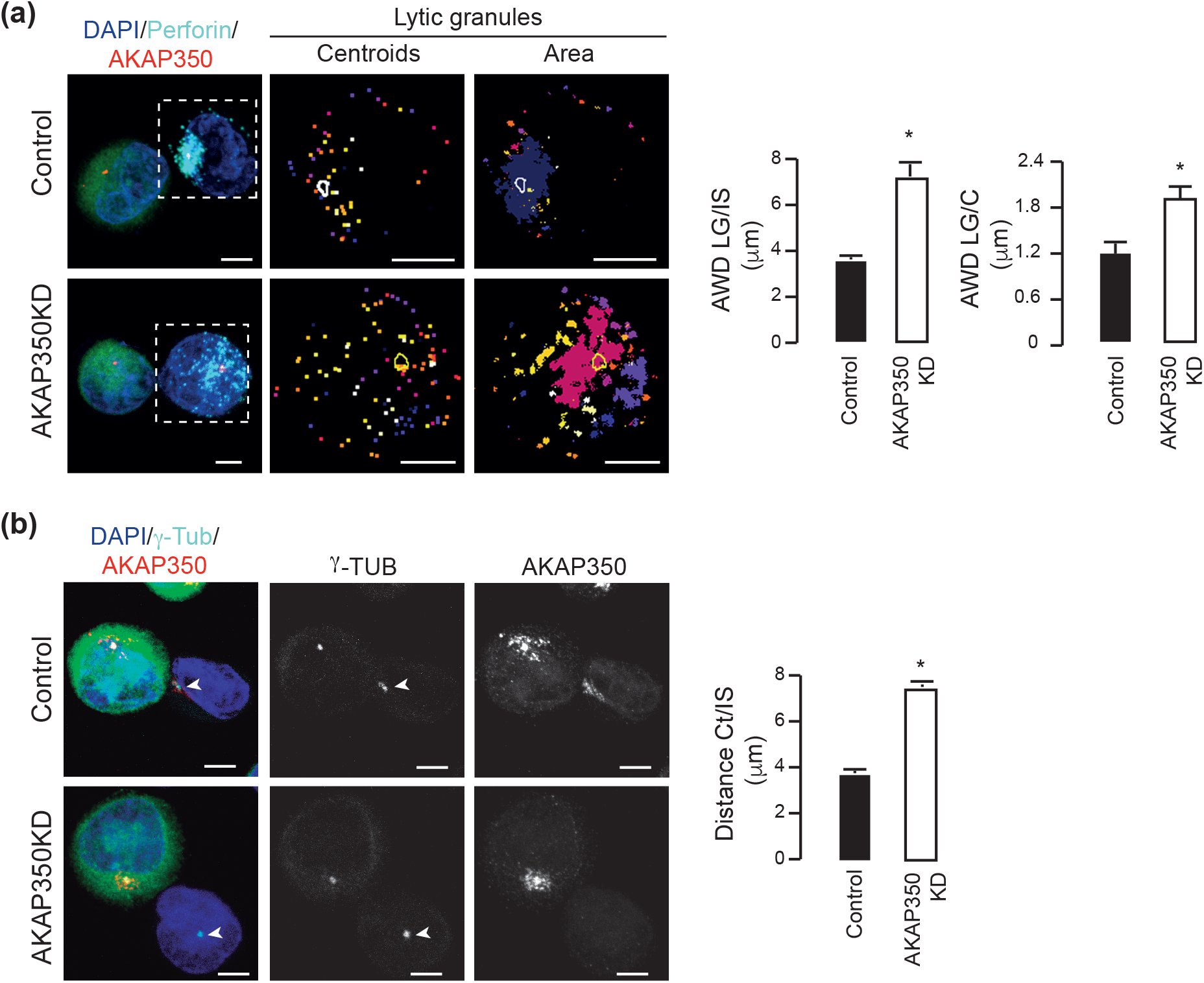
AKAP350 is required for lytic granule convergence and centrosome polarization during YTS cells activation. Control and AKAP350KD YTS cells were incubated with CFSE labeled KT86 cells at a 2:1 ratio for 30 min. Cell conjugates were fixed, stained and analyzed by confocal microscopy. **(a)** Merge images show staining for DAPI (blue) and perforin (cyan) and γ-tubulin (red). The second and third columns show outputs for the lytic granules centroids (second column) or the lytic granules area (third column), corresponding to the boxed cell from the first column, obtained with a specific Image J tool. The localization of the centrosome, as delimited with an automatic selection tool, is outlined in white. Bars represents the mean AWD from the lytic granules to the IS or to the centrosome (C). **(b)** Merge images show staining for DAPI (blue), AKAP350 (red) and γ-tubulin (cyan). White arrows indicate the position of γ-tubulin-labeled centrosomes in YTS cells. Bars represent the mean distance from the centrosome to the IS. Results are representative of four (a) or five (b) experiments. At least 15 conjugates were analyzed for each experiment. *p<0.05. Scale bars, 5 μm

### 3.2 The decrease in AKAP350 expression impairs LFA-1 reorganization at the IS in activated NK cells

Analysis of YTS:KT86 conjugates morphology indicated that the surface of interaction between both cells was reduced in AKAP350KD cells, which was verified by a decrease in the IS area to 50% of control levels (p<0.01). Consistently, both LFA-1 positive area at the IS and LFA-1 average intensity at this area were decreased in AKAP350KD cells, which indicated that LFA-1 accumulation at the IS was impaired in AKAP350KD cells (Figure 3a). In addition, the analysis of the 3D organization of the IS indicated that LFA-1 clustered at the pSMAC in control cells, whereas it was homogeneously distributed in AKAP350KD cells (Figure 3b). It is noteworthy that both total and plasma membrane-associated levels of LFA-1 were preserved in AKAP350KD cells (Supplementary Figure S3), what is in line with our results showing that the decrease in AKAP350 expression did not elicit any evident defect on YTS conjugation to KT86 cells (Supplementary Figure S2).

**Figure 3.**
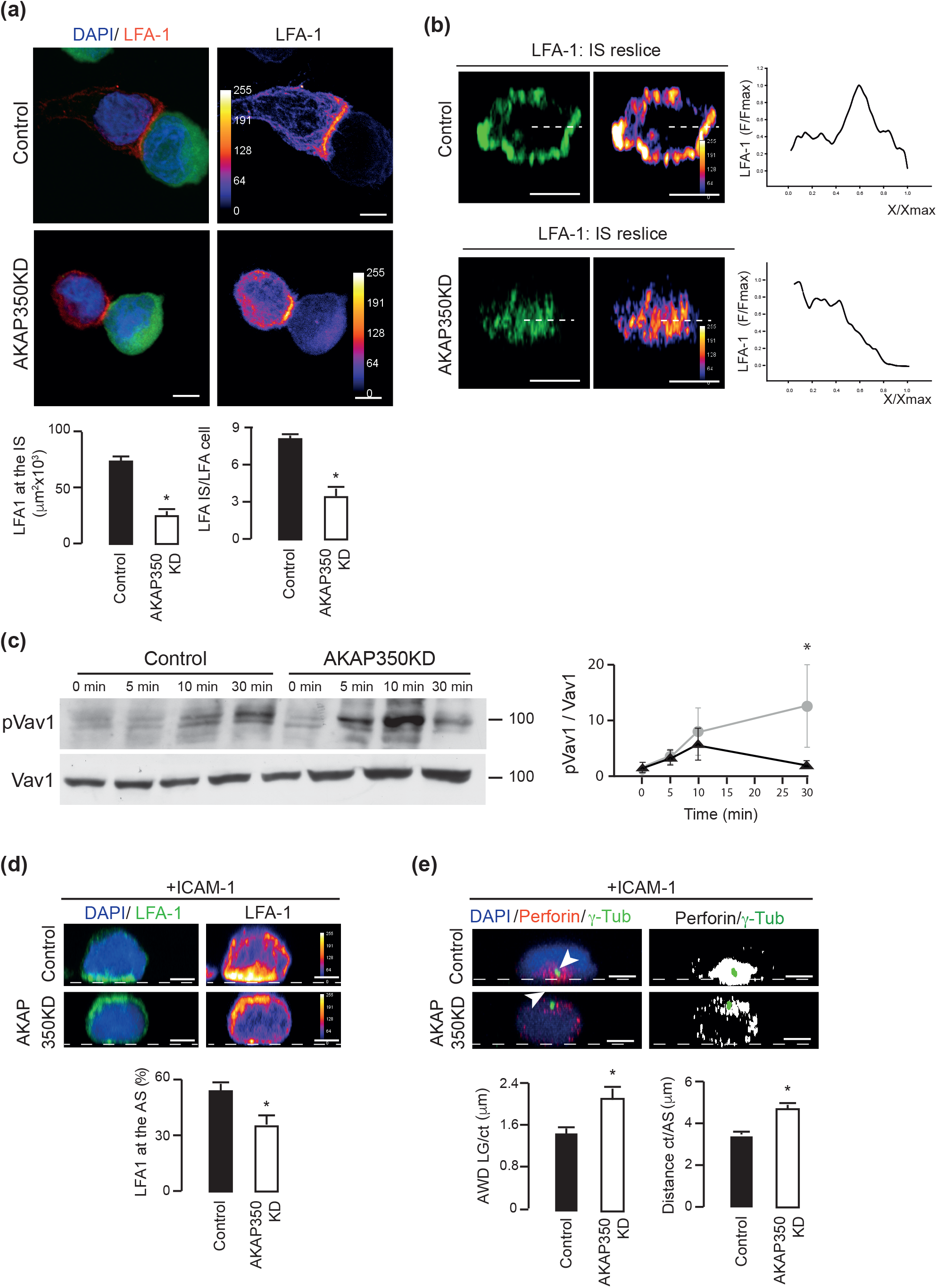
Reduction of AKAP350 expression levels results in defective LFA-1 organization at the IS. **(a)** Control and AKAP350KD YTS cells were incubated with CFSE labeled KT86 cells at a 2:1 ratio for 30 min. Merge images show YTS:KT86 conjugates staining for LFA-1 (red) and DAPI (blue). Bars represent the mean area positive for LFA-1 at the IS and the average intensity of LFA-1 fluorescence at the IS relative to the average intensity of LFA-1 fluorescence at the cell for four experiments. At least 15 conjugates were analyzed for each experiment. Error bars represent SEM. **(b)** Images show reconstructions of the *x,z* reslice of the IS region corresponding to control and AKAP350KD YTS cells. Dashed lines denote the radius of the IS where LFA-1 intensity of fluorescence was quantified. Charts represent the intensity profile of fluorescence, representative of at least 10 IS analyzed in three experiments. **(c)** Western blot analysis of phosphorylated Vav1 (pVav1, upper blot) and total Vav1 (lower blot) levels during YTS activation with KT86 cells, at the indicated periods. The curve chart represents media levels of pVav1 relative to total Vav1 for four experiments. **(d,e)** Control and AKAP350KD YTS cells were activated using coverslips coated with ICAM-1. **(d)** Merge images show LFA-1 (green) and DAPI (blue) staining. White dashed lines indicate the cell surface in contact with the coverslip. Bars represent the mean fraction of LFA-1 fluorescence located at the cell surface in contact with the ICAM-1 coated coverslip (activated surface, AS), expressed as the percentage of total cell fluorescence, representative of three experiments. At least 30 cells were analyzed for each experiment. **(e)** Merge images show γ-tubulin (green), perforin (red) and DAPI (blue) staining. The second column shows the binarized image of the perforin channel, with the ROI corresponding to the centrosome indicated in green. Bars represent the mean AWD between the lytic granules and the centrosome (ct) (first chart) or between the centrosome and the ICAM-1 coated surface (second chart), for at least 15 cells, representative of three experiments. Scale bars, 5 μm (a,d,e) or 2.5 μm (b). *p<0.05

In NK cells, engagement of LFA-1 by ICAM-1 generates early activation signaling by inducing Vav1 activation (Riteau et al., 2003). Vav1 is a nucleotide exchange factor (GEF) for Rac1, which plays an essential role in NK cell signaling pathways that leads to lytic granules polarization and release (Watz and Long, 2010). To examine the potential role of AKAP350 in early events that affect NK signaling, we assessed Vav1 activation during NK:target cell conjugate formation. Time-course activation experiments revealed that Vav-1 phosphorylation was not affected during early activation (5 min and 10 min), but significantly diminished at longer periods (Figure 3c). These results indicate that the initial stages of NK activation were preserved in AKAP350KD cells, whereas there were later events that were impaired.

Differently from T cells, in the absence of inhibitory signaling, LFA-1 activation is enough to induce IS maturation (Barber et al., 2004). Interestingly, when YTS cells were specifically activated with the LFA-1 ligand ICAM-1, inhibition of LFA-1 reorganization was also observed in AKAP350KD cells (Figure 3d), thus suggesting a direct effect of AKAP350 on LFA-1 reorganization. In order to evaluate if the inhibition of lytic IS maturation in AKAP350KD cells (Figure 2) was also elicited when YTS cells were exclusively activated via LFA-1, we further analyzed lytic granule convergence and centrosome translocation towards the activated surface in cells specifically activated with ICAM-1. Our results showed that both processes were inhibited in AKAP350KD cells (Figure 3e).

Altogether, these results indicate that AKAP350 is directly involved in LFA-1 recruitment to the lytic IS in NK cells, and provide evidence for that mechanism being responsible for the defective IS maturation and decreased lytic activity observed in AKAP350KD cells.

### 3.3 NK cells express an intracellular pool of LFA-1 that polarizes to the IS in an AKAP350 dependent manner

The mechanisms underlying LFA-1 accumulation at the IS in NK cells have not been clarified so far. Previous studies in naive CD8+ T cells revealed the presence of an intracellular pool of LFA-1 that translocate to the IS upon T cell activation (Capece et al., 2017). To our knowledge, there was no data regarding the presence of a similar pool of LFA-1 in NK cells. Our analysis of permeabilized YTS cells indicated the presence of cytosolic vesicles containing LFA-1, which were polarized towards the IS in control, but not in AKAP350KD activated cells (Figure 4a). Similarly to what was described in T cells, LFA-1 vesicles partially colocalized with Rab11, indicating partial association with the recycling endosomes (Figure 4b). More interestingly, we found that intracellular LFA-1 also colocalized with GM130 (Figure 4c), which suggested the participation of the Golgi apparatus in LFA-1 intracellular trafficking. LFA-1 colocalization with GM130 or with Rab11 was not affected by the decrease in AKAP350KD expression (Supplementary Figure S4).

**Figure 4.**
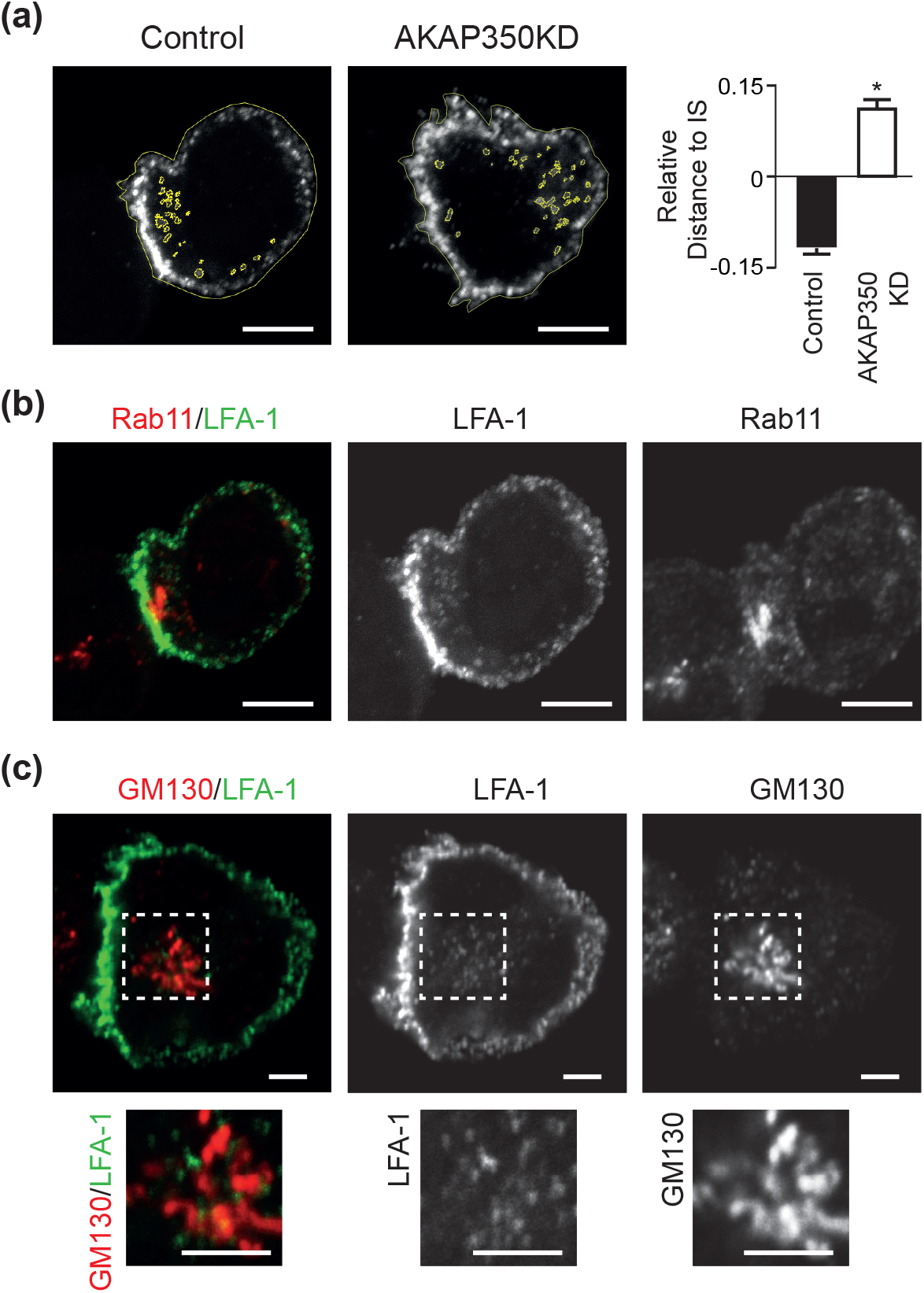
LFA-1 is expressed intracellularly and redistributes to the IS in an AKAP350-dependent manner in YTS cells. Control and AKAP350KD YTS cells were incubated with KT86 cells at a 2:1 ratio for 30 min. **(a)** Images show LFA-1 staining (grey scale) and the ROIs delimiting YTS cells and the individual LFA-1 vesicles in yellow. Bars represent the mean difference between the AWD from LFA-1 vesicles to the IS and the distance from the cell centroid to the IS, relativized to the latter, representative of three experiments. At least 10 conjugates were analyzed for each experiment. **(b,c)** Merge images show staining for LFA-1 (green) and Rab11 (red, b) or GM130 (red, c). **(c)** Amplifications of the boxed areas are shown below each image. Scale bars, 5 μm (a,b) or 2.5 μm (c). *p<0.05

To extend those results to *ex vivo* NK cells, LFA-1 localization was studied in human NK cells prepared from healthy donors. Like YTS cells, *ex vivo* NK cells showed an intracellular pool of LFA-1 which localized in the Golgi apparatus region (Figure 5a). Similarly to YTS cells, ex vivo NK cells transfected with lentiviral particles for expression of AKAP350 shRNAs had a significant decrease in AKAP350 expression (−70%, Figure 5b). In conjugates formed between *ex vivo* NK cells and susceptible target cells, intracellular vesicles containing LFA-1 polarized towards the IS in control, but not in AKAP350KD cells (Figure 5c). Furthermore, LFA-1 localization at the IS was significantly decreased in *ex vivo* NK AKAP350KD cells (Figure 5c).

**Figure 5.**
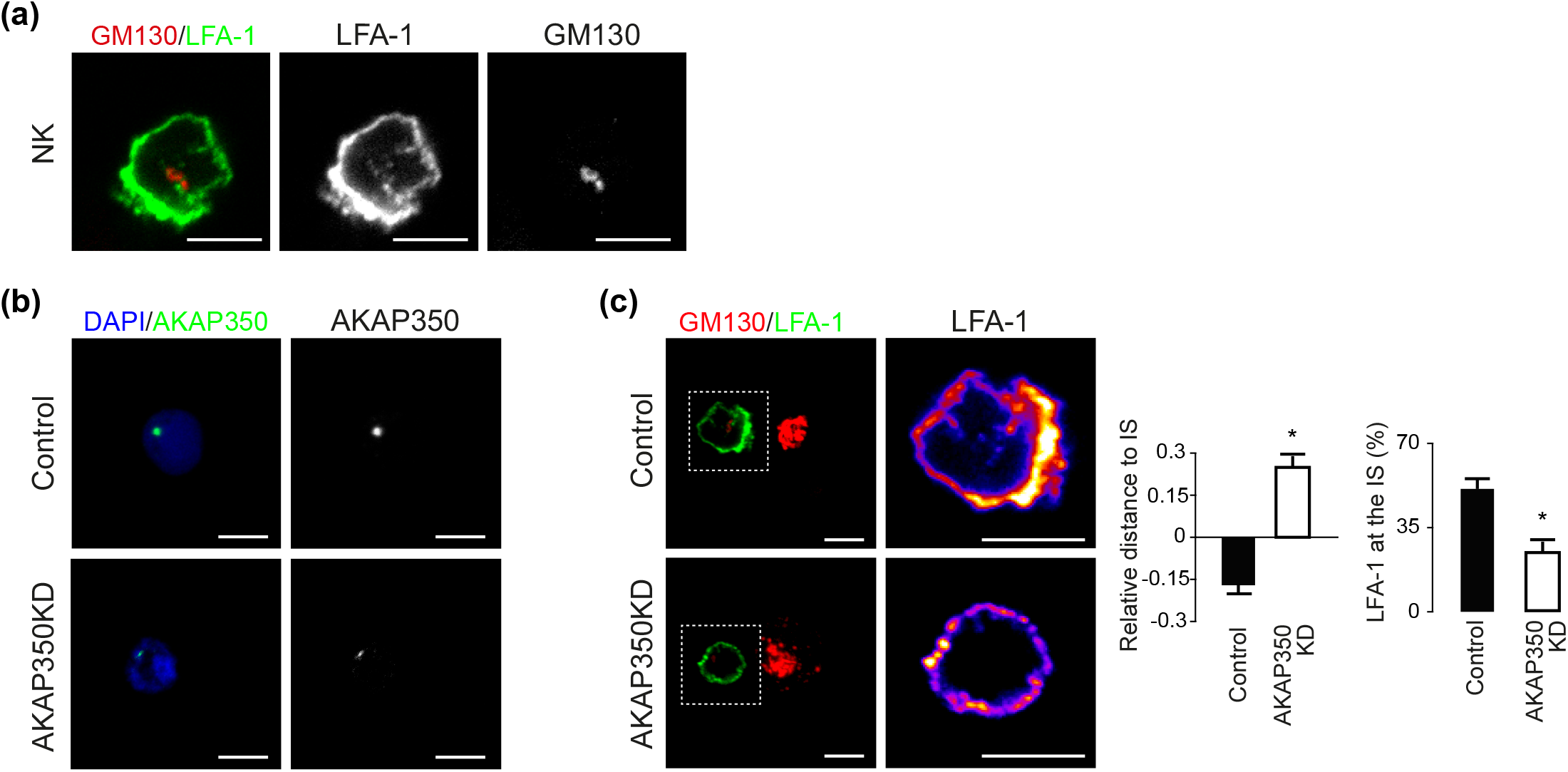
LFA-1 is expressed intracellularly and redistributes to the IS in an AKAP350-dependent manner in *ex vivo* NK cells. NK cells were prepared from blood samples from volunteer donors as described in Materials and methods. **(a)** NK cells were incubated with KT86 cells at a 2:1 ratio for 30 min. Merge image shows staining for LFA-1 (green) and GM130 (red). **(b,c)** Cells were transduced with lentiviral particles codifying for control or AKAP350 specific shRNAs. Transduced cells were selected with puromycin for 96 h. **(b)** Merge images show staining for AKAP350 (green) and nuclei (blue) in control and AKAP350KD cells. **(c)** Control and AKAP350KD NK cells were incubated with KT86 cells and processed as in (a). Merge images show staining for LFA-1 (green) and GM130 (red). For better visualization of LFA-1 subcellular distribution, an amplification of the LFA-1 channel of the boxed area is shown in pseudocolor. Bar charts represent the average relative distance from LFA-1 vesicles to the IS, calculated as in Figure 4a, (left), and the mean fraction of LFA-1 fluorescence located at the IS, expressed as percentage of total cell fluorescence for the LFA-1 channel (right), representative of three independent ex vivo NK cell preparations. *p<0.05

### 3.4 AKAP350 association to the GA is essential for NK-IS maturation

Considering the partial colocalization of intracellular LFA-1 with the Golgi apparatus, we examined the relevance of Golgi associated AKAP350 in the maturation of NK lytic IS. We first verified that AKAP350 had a prominent localization at the Golgi apparatus in activated NK cells (Supplementary Figure S5). Analysis of Golgi and centrosomal localization of AKAP350 in those cells further suggest that NK activation induced partial translocation of AKAP350 from the centrosome towards the Golgi apparatus (Supplementary Figure S5). In order to analyze the relevance of Golgi associated AKAP350 in LFA-1 organization in activated YTS cells, we prepared YTS cells with stable expression of AKAP350(1-1229) (AKAP350GBD) domain, which acts as a dominant negative construct that displaces the protein from the Golgi apparatus (Hurtado et al., 2011). The analysis of AKAP350 colocalization with GM130 confirmed that AKAP350 localization at the Golgi apparatus was significantly decreased in AKAP350GBD cells (Supplementary Figure S6). The analysis of the intracellular pool of LFA-1 showed that, whereas LFA-1 colocalization with GM130 was preserved in AKAP350GBD cells (Figure 6a), its distribution towards the IS was impaired in those cells (Figure 6b). Concomitantly, LFA-1 accumulation and organization at the IS was compromised in AKAP350GBD cells (Figure 6c, d). Overall, this set of data reveal the participation of Golgi-associated AKAP350 in LFA-1 relocalization to the IS.

**Figure 6.**
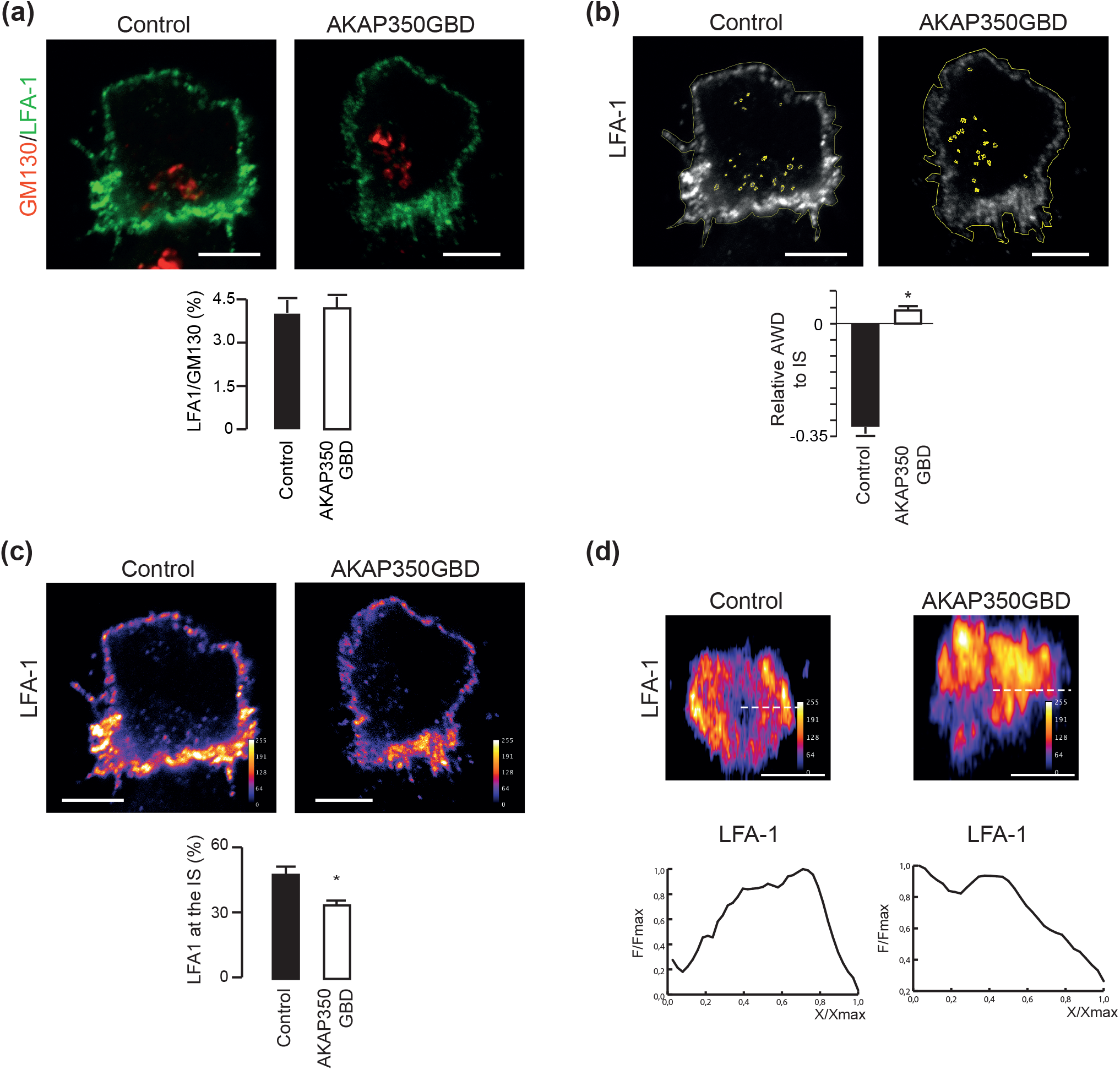
AKAP350 localization at the Golgi apparatus is required for IS maturation. Control and AKAP350GBD YTS cells were incubated with KT86 cells. **(a)** Merge images show staining for LFA-1 (green) and GM130 (red). Bars represent the mean fraction of LFA-1 that colocalized with GM130 for 10 different cells. **(b)** Images show LFA-1 staining (grey scale) and the ROIs delimiting the cells and the individual LFA-1 vesicles (yellow). Bars represent the mean difference between the AWD from LFA-1 vesicles to the IS and the distance from the cell centroid to the IS, relativized to the latter, representative of three independent experiments. At least 10 conjugates were analyzed for each experiment. **(c)** Images show LFA-1 staining in pseudocolor. Bars represent the mean fraction of LFA-1 fluorescence located at the IS, expressed as a percentage of total cell fluorescence. More than 30 conjugates from three different experiments were analyzed. **(d)** Images show reconstructions of the *x,z* reslice at the IS region for LFA-1 staining, corresponding to control and AKAP350GBD YTS cells. Dashed lines denote the radius of the IS where LFA-1 intensity of fluorescence was quantified. Charts represent the intensity profile of fluorescence, representative of at least 10 IS. Scale bars, 5 μm (a-c) or 2.5 μm (d). *p<0.05

### 3.5 Disruption of Golgi integrity by BFA leads to defective LFA-1 organization at the IS

To further assess the implication of the Golgi apparatus in LFA-1 recruitment to the IS in activated cells, we analyzed LFA-1 localization at the IS in NK cells that were pre-treated with Brefeldin A (BFA), a pharmacologic inhibitor of ADP-ribosylation factors that leads to the collapse of the Golgi and its redistribution into the ER (Chardin and McCormick, 1999). Analysis of LFA-1 distribution at the IS showed that BFA-treatment inhibited LFA-1 localization at the IS (Figure 7a) and organization at the pSMAC (Figure 7b).

**Figure 7.**
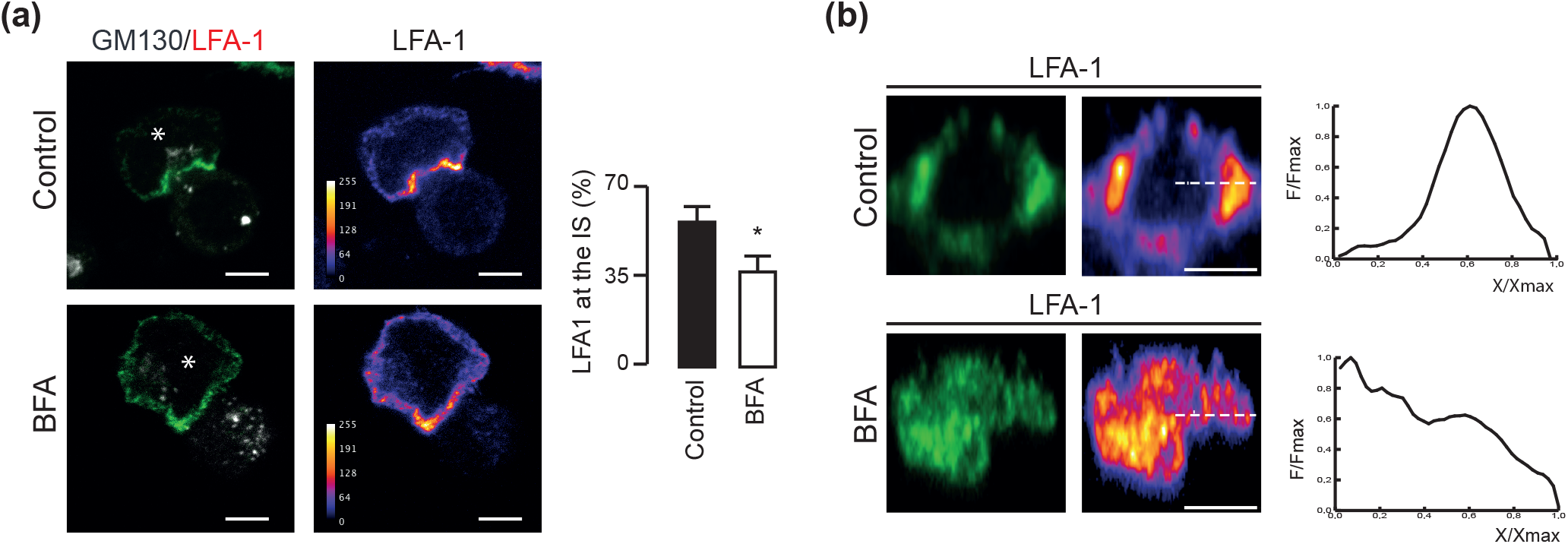
LFA localization at the IS is dependent on the integrity of the Golgi apparatus. YTS cells were treated with brefeldin A (BFA) as described in Material and methods. Control or BFA treated YTS cells were incubated with KT86 cells. **(a)** Merge images show staining for LFA-1 (green) and GM130 (gray). Asterisks indicate YTS cells. Bars represent the mean fraction of LFA-1 fluorescence located at the IS, expressed as percentage of total cell fluorescence for the LFA-1 channel, representative of three independent experiments. At least 8 conjugates were analyzed for each experiment. **(b)** Images show reconstructions of the *x,z* reslice of the IS region corresponding to control and BFA treated YTS cells. Dashed lines denote the radius of the IS where LFA-1 intensity of fluorescence was quantified. Charts represent the intensity profile of fluorescence, representative of at least 10 IS. Scale bars, 5 μm (a) 2.5 μm (b). *p<0.05

### 3.6 Microtubule cytoskeleton integrity is necessary for LFA-1 organization at the IS

To further verify that vesicle trafficking was required for LFA-1 organization at NK-IS, we analyzed if microtubule integrity was required for LFA-1 accumulation at the IS. We found that both cold-induced microtubule depolymerization in the presence of Nocodazole or suppression of microtubule dynamic instability with Taxol inhibited LFA-1 accumulation and distribution at the IS (Figure 8). In the same conditions, the reorganization of the actin cytoskeleton at the IS was preserved.

**Figure 8.**
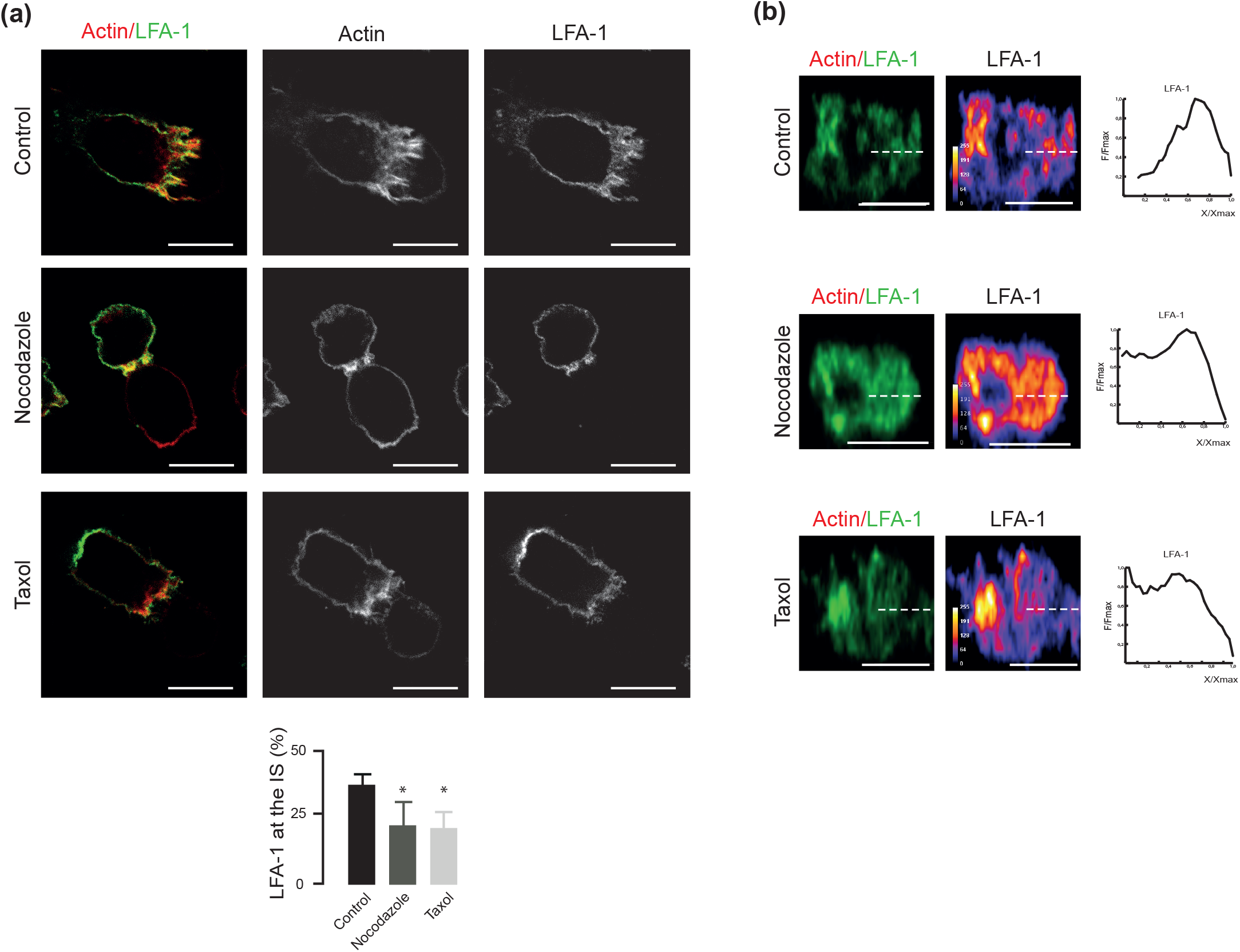
Microtubule-integrity is required for LFA-1 organization at the IS. Control YTS cells or YTS cells treated with nocodazole or taxol were mixed with KT86 cells at a 2:1 ratio. **(a)** Merge images show cell conjugates staining for actin (red) and LFA-1 (green). Bars represent the mean fraction of LFA-1 fluorescence located at the IS, expressed as a percentage of total cell fluorescence. More than 30 conjugates from three different experiments were analyzed. **(b)** Images show reconstructions of the *x,z* reslices of the IS region corresponding to control and nocodazole or taxol treated YTS cells. Dashed lines denote the radius of the IS where LFA-1 intensity of fluorescence was quantified. Charts represent the intensity profile of fluorescence quantified for the LFA-1 channel, representative of at least 10 IS analyzed in three independent experiments. Scale bars, 5 μm (a) 2.5 μm (b).. *p<0.05.

## 4 DISCUSSION

In NK cells, besides its canonical role in the consolidation of cell adhesion to target cells, LFA-1 clustering at the IS plays also a central function in the activation of signaling pathways that lead to lytic granule-translocation towards the IS (Barber et al., 2004; Bryceson et al., 2005). Even though many studies have contributed to clarify the signaling pathways that promote LFA-1 conformational activation during NK interaction with sensitive target cells, the mechanisms leading to LFA-1 accumulation at the IS at these cells have not been elucidated so far. In the present study, we demonstrate that AKAP350 participates in NK cytolytic activity by facilitating LFA-1 recruitment to the lytic IS. Regarding the mechanisms involved, our study reveals the presence of an intracellular pool of LFA-1 that polarizes to the IS in an AKAP350 manner, and provides evidence supporting a prominent role for the Golgi apparatus in LFA-1 organization at the IS.

Our findings show that the decrease in AKAP350 expression leads to decreased NK cytolytic activity, which is associated with defective LFA-1 recruitment to the IS. LFA-1 contribution to early signaling for cytotoxicity in NK cells is very different from integrin-mediated activation of T cells. In contrast with cytotoxic T cells, binding of LFA-1 to ICAM-1 does not need further inside out signaling and is sufficient to promote polarization of cytolytic granules in NK cells (Barber et al., 2004; Bryceson et al., 2005). Analysis of lytic granule polarization towards the activated surface in cells that were exclusively activated with an LFA-1 ligand, ICAM-1, indicates that LFA-1 was, at least in part, primary responsible for the defective NK cytolytic activity in AKAP350KD cells. Both LFA-1 expression at the plasma membrane in resting cells and NK:target cell conjugate formation were preserved in AKAP350KD cells. Furthermore, Vav1 activation, which is an early signaling event where LFA-1 activation converges with other activation signaling pathways (Watz and Long, 2010), was preserved up to 10 min of NK activation, indicating that the initial phase of NK activation was not affected in AKAP350KD cells. In T cells, in contrast, the decrease in AKAP350 expression inhibits Vav1 phosphorylation and LFA-1 signaling 5 min after T cell-APC conjugation, what occurs concurrently with decreased T cell receptor activation (Robles Valero et al., 2010). In those cells, LFA-1 activation depends strongly on T cell receptor signaling, involving signaling molecules that are not involved in LFA-1 activation in NK cells (Watz and Long, 2010). Therefore, initial LFA-1 signaling might not require AKAP350 expression in NK cells, whereas LFA-1 deficient early signaling in AKAP350KD T cells might be secondary to the decreased T cell receptor activation. On the other hand, 30 min after conjugate formation Vav1 activation was significantly decreased in AKAP350KD cells, which is in accordance with the decreased LFA-1 accumulation at the IS observed at this period of activation (Figure 3). Altogether, those results indicate that AKAP350 participates in events primarily involving LFA-1 organization at NK-IS, which occur downstream initial NK cell conjugate formation and activation.

Most of the information regarding LFA-1 redistribution at the cell surface originates in studies performed in T cells, which, as we mentioned above, have prominent differences in regards to LFA-1 activation. The classical model proposes that conformational-ligand induced regulation of LFA-1 lateral motility can account for actin-dependent LFA-1 accumulation at the IS (Cairo et al., 2006). More recently, Kim and col. identified a novel intracellular LFA-1 pool, which actively redistributes to T cell:target cell interaction surface upon antigen stimulation (Capece et al., 2017). In the same line of evidence, previous studies performed in migrating T cells indicated that LFA-1-dependent migration decreases when intracellular vesicle transport is blocked, thus revealing the relevance of an intracellular pool of LFA-1 in the regulation of LFA-1 organization at the plasma membrane in those cells (Stanley et al., 2012). The characterization of that intracellular pool in T cells showed that it associates with early endosomes and with recycling endosomes (Ong et al., 2014; Samuelsson et al., 2017). Our results in YTS and in ex vivo NK cells demonstrate the presence of an intracellular pool of LFA-1 that polarizes towards the IS in control but not in AKAP350 KD cells. Interestingly, that pool of LFA-1 vesicles shows partial association with the Golgi apparatus, suggesting that this organelle could regulate LFA-1 trafficking to the IS in NK cells. Even though it is well established that the Golgi apparatus translocates to the IS during NK lytic response (Kupfer et al., 1983), the role of this organelle in the development of NK lytic IS has not been examined so far. Interestingly, our results show that BFA treatment impairs LFA-1 accumulation at the IS, supporting our hypothesis that the Golgi apparatus regulates LFA-1 traffic to the IS and providing evidence, for the first time, for the participation of the Golgi apparatus in LFA-1 organization at the IS in NK cells.

Previous studies in T cells suggest that AKAP350 participation in T cell activation and migration is attributable to centrosome-associated AKAP350 (El Din El Homasany et al., 2005; Robles-Valero et al., 2010). As mentioned above, AKAP350 participates in Golgi apparatus organization (Larocca et al., 2004). Furthermore, in epithelial derived migratory cells AKAP350 localization at the Golgi apparatus is necessary for directional vesicular traffic towards the leading edge (Hurtado et al., 2011). Considering the results discussed above, we analyzed if there was a specific role for Golgi-associated AKAP350 in IS maturation in NK cells. We found that AKAP350 delocalization from the Golgi apparatus inhibits LFA-1 accumulation and clustering at the IS, ratifying the role of the Golgi apparatus in IS maturation and underscoring the relevance of AKAP350 location at this organelle for the development of NK lytic IS. We speculate that the deficient polarization of the Golgi apparatus towards the IS observed in AKAP350KD and in AKAP350GBD NK cells (Figure 5–6) could be the reason for the impaired LFA-1 trafficking towards this structure in those cells. The characterization of the mechanisms involved in AKAP350-dependent Golgi reorganization towards the NK-IS will be subject of future studies.

We found that levels of nocodazole that actually inhibits NK cytotoxicity (Katz et al., 1982), induce a defect in LFA-1 organization at the IS. Orange et al have previously demonstrated that cell-surface receptor accumulation at the activating NK-IS was not disrupted with colchicine (2003). Colchicine and nocodazole inhibit microtubule polymerization by similar mechanisms (Florian and Mitchison, 2016). In our study, we additionally induced microtubule depolymerization by ice incubation, which may explain the different outcomes between both experimental settings. The fact that levels of the microtubule stabilizing agent Taxol, which impede microtubule growing and shortening events (Derry et al., 1995), also inhibit LFA-1 reorganization at the IS provides further evidence that microtubule dynamics is essential for this process. It is noteworthy that, in our settings, neither Nocodazole nor Taxol treatment affected actin organization at the IS, thus ruling out that the inhibition in LFA-1 organization at the IS could be secondary to microtubule effects on actin cytoskeleton. Altogether, those results further support our hypothesis that intracellular trafficking is required for LFA-1 reorganization at the IS in NK cells.

In conclusion, the present study reveals a novel, AKAP350-dependent pathway that participates in the regulation of NK cytotoxic function, providing evidence for the involvement of a Golgi/AKAP350 dependent-polarized trafficking of LFA-1 in the promotion of LFA-1 organization at the IS during NK cell-activation. The fact that disruption of the Golgi integrity leads to defective LFA-1 organization at the IS indicate that that is a primary pathway involved in the regulation of NK effector function. The significance of a similar mechanism in the development of different types of IS or in immune cells migration represents an interesting subject for further investigation.

## ETHICS APPROVAL AND CONSENT TO PARTICIPATE

All human samples were obtained after informed patient consent and were used with approval of the institutional internal review board for the protection of human subjects of the Hospital Provincial del Centenario (Rosario, Argentina).

## AUTHORS CONSENT FOR PUBLICATION

All the authors give their consent for publication.

## AVAILABILITY OF DATA AND MATERIAL

The datasets and material generated and/or analyzed during the current study are available from the corresponding author on reasonable request.

## ACKNOWLEDGMENTS

We thank Dr. Norberto Zwirner for providing us with reagents and advice for NK cell purification and Dr. Andrés Zucchetti for his critical discussion of the manuscript.

