## Supplementary methods and figures for "IDENTIFICATION OF A NOVEL MECHANISM FOR LFA-1 ORGANIZATION DURING NK CYTOLYTIC RESPONSE"

SUPPLEMENTARY METHODS (SM)

SM1 Preparation of lentiviral particles and generation of AKAP350KD cells

Constructs were made by annealing and ligating oligonucleotides targeting two specific AKAP350 sequences (shRNA1, 5′-CCCAGCTCACTGCTAATTT-3′; shRNA4, 5′-GCAAGAACTAGAACGAGAA-3′) or a scrambled control into the AgeI and EcoRI cloning sites of pLKO.1-puro vector (details at http://www.addgene.org). These constructs were sequenced and used to co-transfect human embryonic kidney 293 FT cells with Virapower lentiviral packaging mix (Invitrogen, Carlsbad, CA). The next day, transfection complexes were removed, and cells were allowed to produce virus for 24 h. Media containing viruses were collected and used to directly transduce YTS cells overnight. The cells were then allowed to recover for 24 h and subjected to puromycin selection (2 μg/ml) for 2 weeks. Silencing was confirmed by western blotting and immunofluorescence analyses.

SM2 *Ex vivo* NK cell purification

In order to separate red blood cells from leukocytes, 20 ml of whole blood sample was centrifuged at 900g for 15 minutes and the buffy coat was obtained. The buffy coat was resuspended in an equal volume of PBS and the Peripheral Blood Mononuclear Cells (PBMC) fraction was obtained by centrifugation on a Ficoll cushion for 30 minutes at 600g. The PBMC were washed twice with physiological solution and centrifuged for 15 minutes at 400g. The PBMC were resuspended in 500 uL of whole blood and incubated for 20 minutes at room temperature with RosetteSep Human NK cells Enrichment cocktail (StemCell # 15025) to purify NK cells by negative selection. After the incubation period, a  volume of physiological solution supplemented with 2% of FBS was added and the NK cells enriched fraction was obtained by centrifugation on a Ficoll cushion for 20 minutes at 400g. Purified NK cells were washed twice in physiological solution supplemented with 2% FBS for 10 minutes at 400g, resuspended and maintained in the same medium used for YTS cells, supplemented with IL-2 500 UI/ml at 37°C and 5% $CO_2$ atmosphere.

SM3 FACS-based conjugation assay

YTS labeling reaction was stopped by the addition of an equal volume of fetal bovine serum, followed by a 5 min incubation at room temperature. After two washes with PBS, the YTS-labeled cells were resuspended in RPMI complete medium. Cells were combined at an effector to target ratio of 2:1 or 10:1 and incubated at 37º C for different periods. The samples were gently vortexed for 3 s and immediately fixed with 2% paraformaldehyde (PFA). Samples were run in triplicates and 50000 events were counted for each replicate. The frequency of double positive events was determined within the total cell population using FlowJo software. The following gating strategy was used: First, cells were gated to exclude debris. Compensation adjustments were made on the gated population using single-positive cells stained for either Cell TrackerTM Deep Red or CFSE. Gates were set to differentiate between the double positives, represented in G2, from the single positives and double negatives. The percentage of conjugated cells was calculated as follows: conjugated cells (%) = [Double positive cells / total cells] x 100.

SM4 Immunofluorescence

In experiments evaluating NK cell activation by means of specific activating receptors, non adherent cells were washed and cells adhering to the slide were fixed with 4% PFA at room temperature or methanol at -20°C. In every case, fixed cells were blocked with 1% bovine serum albumin/PBS, pH 7.4, for 10 min. When indicated, 0.3%Triton X-100 was added to the buffer to enhance cell permeabilization. Then, they were incubated for 2 h with mouse monoclonal antibody anti-LFA-1 (Biolegend 301202), anti-AKAP350 (Schmidt et al, 2001), anti-Perforin

(Santa Cruz SC-136994) or anti-alpha tubulin (Sigma T5168) and rabbit monoclonal antibody anti γ-tubulin (Sigma-T5192), anti-GM130 (Abcam EP892Y) or anti-Rab11 (Abcam ab180504). The coverslips were washed, incubated for 1 h with the secondary antibodies conjugated to Alexa 488, Alexa 560 or Alexa 633 or with phalloidin-Alexa 568 (Molecular probes-A34055, 1:200) for actin staining and with 4′,6-diamidino-2-phenylindole (DAPI) and mounted with ProLong (Invitrogen). Fluorescence was detected using LSM880 confocal with an ObserverZ1 inverted microscope. Serial optical 0.4 µm thick sections were collected in the z-axis. Z-stacks were built, and projections were obtained using ImageJ tools. In preparing the figures, adjustment in brightness and contrast were equally applied to the entire images using Adobe Photoshop software to improve visualization of fluorescence.

SM5 Immunoblotting

Cells were lysed in ice-cold lysis buffer (50 mM Tris-HCl [pH 7.5], 100 mM NaCl, 15 mM EDTA and 1% Triton X-100, with protease inhibitors) and subjected to two freeze–thaw cycles. Lysates were centrifuged at 1000 *g* for 5 min at 4°C, and the clear supernatants were conserved. For all samples, total protein concentrations were measured according to Lowry *et al.* (1951). Samples were heated for 10 min at 90°C in sample buffer (20 mM Tris-HCl, pH 8.5, 1% SDS, 400 µM DTT, 10% glycerol). Samples containing equal amounts of proteins were subjected to SDS polyacrylamide gel electrophoresis (6–10% gradient to verify AKAP350 silencing or 10% otherwise). The proteins in the 6% gel were transferred to nitrocellulose membranes (Amersham Pharmacia Biotech), whereas the 10% gel proteins were transferred to polyvinyl difluoride membranes (Perkin Elmer Life Sciences). Blots were blocked with 5% non-fat dry milk in PBS, 0.3% Tween-20 (PBS-Tween). Nitrocellulose blots were then probed with the monoclonal mouse anti-AKAP350 antibody (1:500), and polyvinyl difluoride membranes were probed with mouse monoclonal antibodies anti-α-tubulin (Sigma-T5168, 1:5000), anti-CIP4 (BD Bioscience 612557), polyclonal rabbit antibody anti-phosphoVAV1 (ThermoFisher PA1-26651, 1:1000) or monoclonal rabbit antibody anti-VAV1 (Cell Signaling 4657S, 1:1000). The blots were washed and incubated with the horseradish peroxidase-conjugated corresponding secondary antibodies, and bands were detected by enhanced chemiluminescence (Pierce, Thermo Scientific). Autoradiographs were obtained by exposing the blots to Kodak XAR film. The bands were quantitated by densitometry using the NIH Image J program.

### Supplementary Figure S1

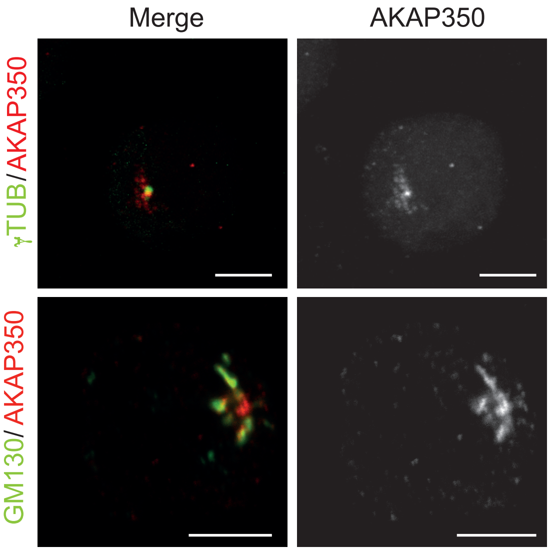


AKAP350 localization in YTS cells. Isolated YTS cells were stained and analyzed by confocal microscopy. Merge images show YTS cells staining for AKAP350 (red) and γ-tubulin or GM130 (green). Scale bars, 5 µm

**Supplementary Figure S2**

**(a)**

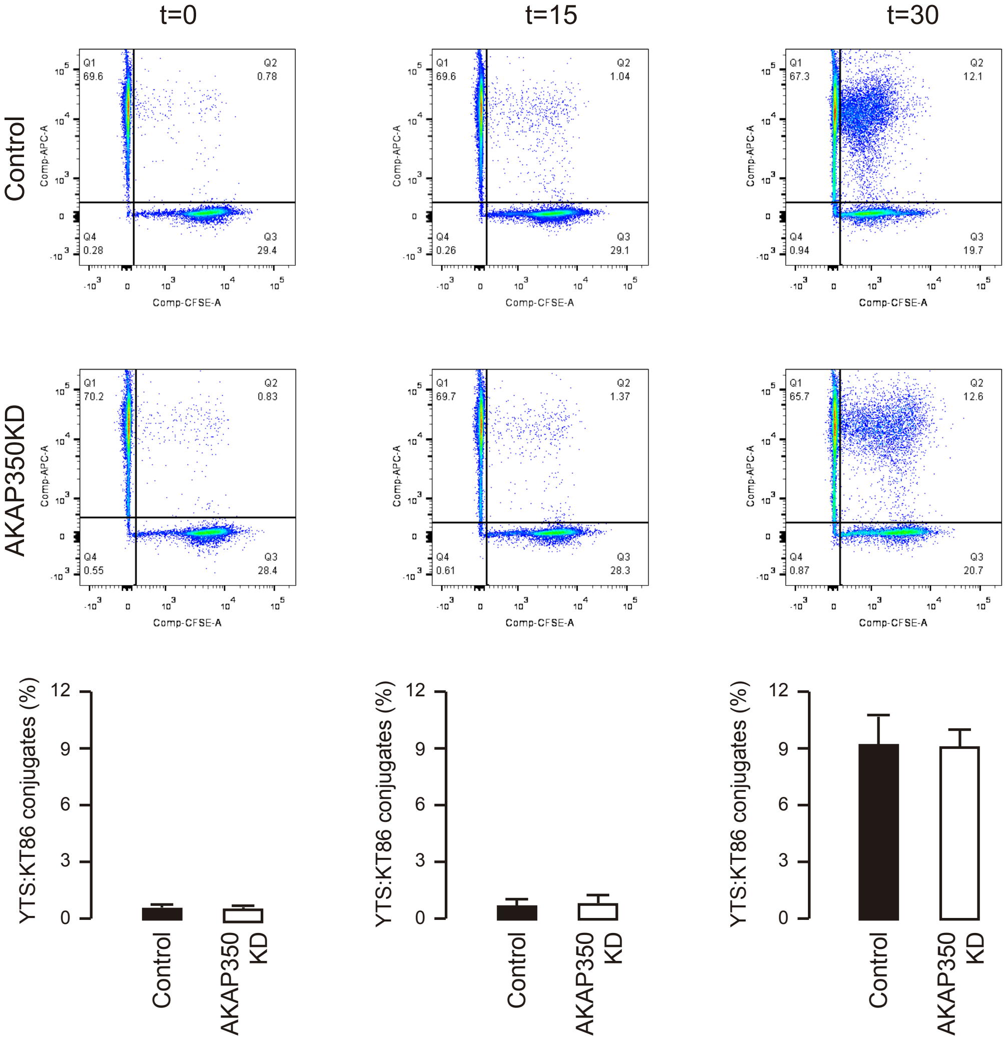


**(b)**

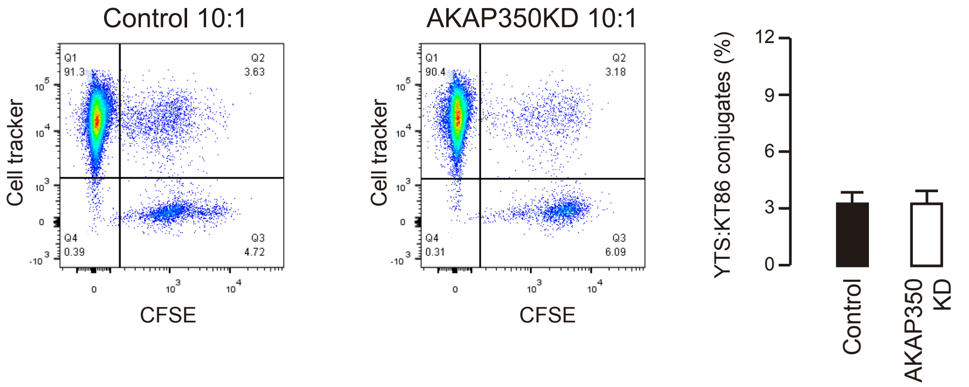


Reduction of AKAP350 expression levels does not affect YTS:KT86 conjugate formation. YTS control and AKAP350KD cells were stained with cell tracker Deep red and mixed with KT86 cells stained with CFSE. Conjugate formation was analyzed by flow cytometry. The conjugates were determined as CFSE/cell tracker double positive events. Effector and target cells were mixed at a 2:1 (**a**) or 10:1 (**b**) ratio and incubated at 37 °C and 5 % $CO_2$ for the indicated periods (**a**) or for 30 min (**b**). Bars represent the mean percentage of doble positive events for three independent experiments. Error bars represent SEM. *$p < 0.05$

**Supplementary Figure 3**

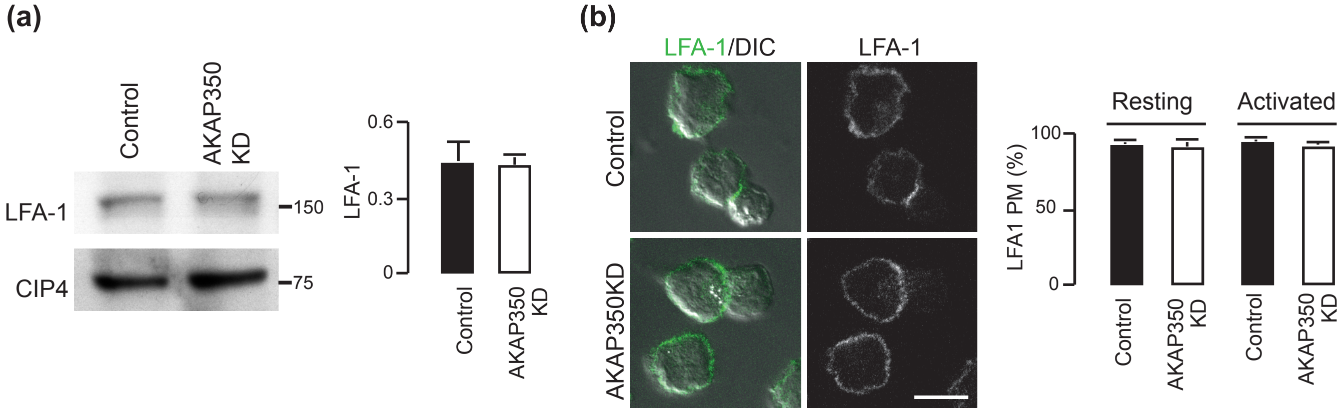


The decrease in AKAP350 expression does not affect LFA-1 expression or localization at the plasma membrane. **a** Western blot analysis of LFA-1 expression in control and AKAP350KD YTS cell lysates. CIP4 was used as loading control. Bars represent the mean density for LFA-1, relativized to the loading control, for three independent experiments. **b** Control and AKAP350KD YTS cells were incubated with KT86 cells. Merge images show staining for LFA-1 in green and the DIC in grey-scale. Bars represent the mean fraction of LFA-1 that localized at the plasma membrane in isolated (resting) YTS cells, and in YTS cells conjugated to KT86 cells (activated) for 10 different cells in each group. Error bars represent SEM. Scale bars, 10 μm

### Supplementary Figure S4

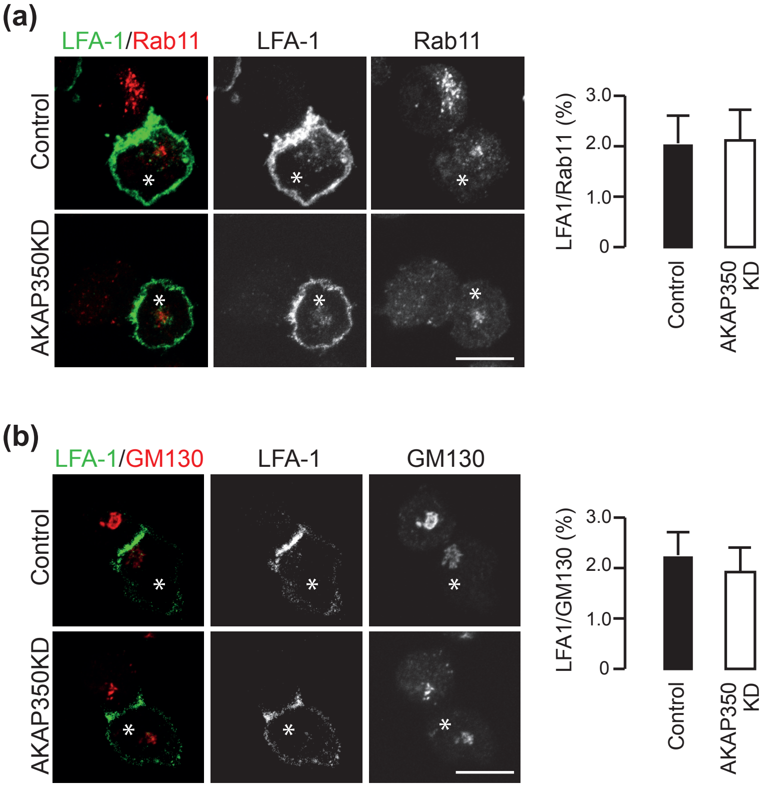


The decrease in AKAP350 expression does not affect LFA-1 localization at the recycling endosomes or at the Golgi apparatus. Control and AKAP350KD YTS cells (asterisk) were incubated with KT86 cells. Merge images show staining for LFA-1 in green, and for Rab11 (**a**) or GM130 (**b**) in red. Bars represent the mean fraction of LFA-1 that colocalized with Rab11 (**a**) or with GM130 (**b**) for 10 different cells. Error bars represent SEM. Scale bars, 10 μm.

### Supplementary Figure S5

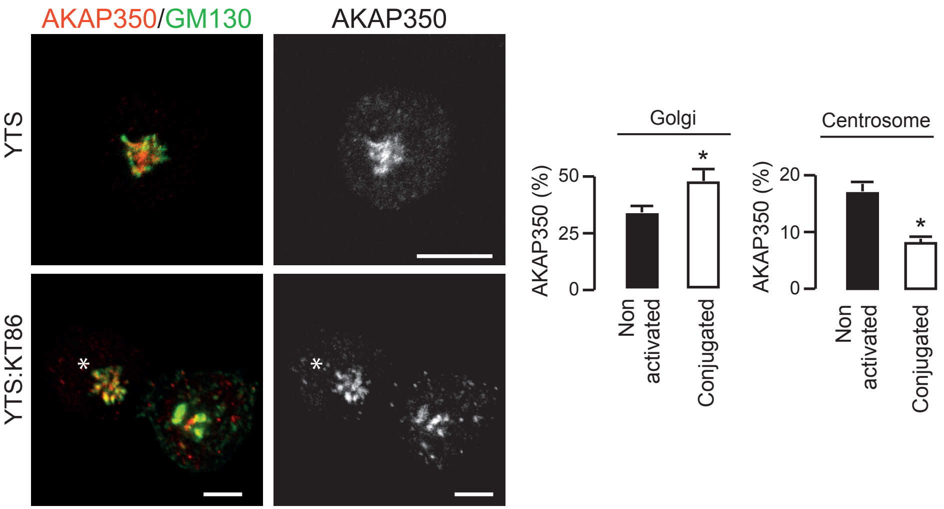


AKAP350 localization in YTS:KT86 conjugates. Merge images show YTS cells (upper row) or YTS:KT86 conjugates (lower row) staining for AKAP350 (red) and GM130 (green). Asterisks denote YTS cells in YTS:KT86 conjugates. Bars represent the mean fraction of AKAP350 fluorescence present at the Golgi apparatus or at the centrosomes, expressed as a percentage of total AKAP350 fluorescence for each cell. Results are representative of three independent experiments. At least 20 cells were analyzed for each experiment. Error bars represent SEM. *$p < 0.05$. Scale bars, 5 μm

### Supplementary Figure S6

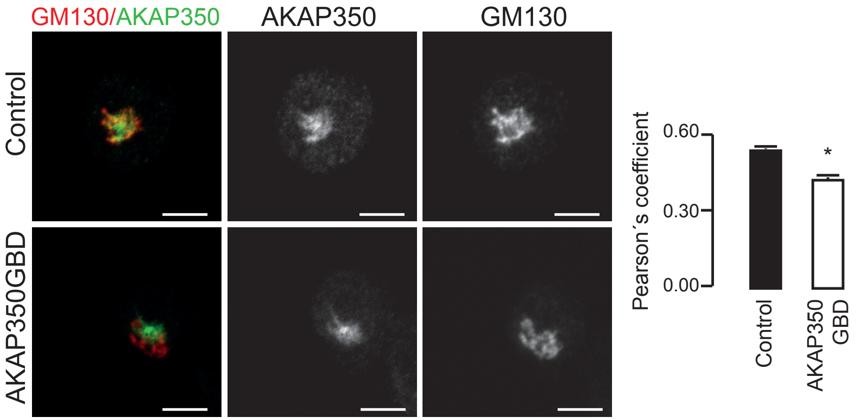


AKAP350 localization at the Golgi apparatus is reduced in AKAP350GBD cells. Merge images show staining for AKAP350 (green) and GM130 (red). Bars represent the mean Pearson correlation coefficient (R) for AKAP350 and GM130 channels, used to estimate AKAP350 colocalization with GM130, for 10 different cells. Error bars represent SEM. Scale bars, 5 µm.*$p<0.05$.
